# Antibiotics effects on the fecal metabolome in preterm infants

**DOI:** 10.1101/2020.06.26.159590

**Authors:** Laura Patton, Nan Li, Timothy J. Garrett, J. Lauren Ruoss, Jordan T. Russell, Diomel de la Cruz, Catalina Bazacliu, Richard A. Polin, Eric W. Triplett, Josef Neu

## Abstract

Within a randomized prospective pilot study of preterm infants born less than 33 weeks gestation, fecal samples were collected weekly and metabolomic analysis was performed. The objective is to evaluate for differences in fecal metabolites in infants exposed to antibiotics vs not exposed to antibiotics in the first 48hours after birth. Significant differences were seen in the antibiotics vs no antibiotics group, including pathways related to vitamin biosynthesis, bile acids, amino acid metabolism and neurotransmitters. Early antibiotic exposure in preterm infants may alter metabolites in the intestinal tract of preterm infants. Broader multi-omic studies that address mechanisms will guide more prudent antibiotic use in this population.

## Introduction

Antibiotics are the most common drugs prescribed to infants in the neonatal intensive care unit (NICU), largely because of the concern for sepsis^[1,2,3]^. The dogma for most neonatologists is that providing pre-emptive routine antibiotics for very low birthweight infants right after birth saves lives by decreasing sepsis without attendant complications. Currently, evidence to support the practice of routinely administering antibiotics and the lack of complications is inadequate. Despite recent initiatives to limit antibiotic use and the low incidence of early onset sepsis (EOS), most infants born less than 33 weeks’ gestational age (GA) are exposed to a course of antibiotics shortly after birth. Recent trials suggest that use of antibiotics in preterm infants have adverse consequences such as an increase in necrotizing enterocolitis, chronic lung disease and death ^[4,5,6]^. Understanding the long-term effects of antibiotics on preterm infants may further guide this practice.

It is well known that intestinal microbial metabolism plays a major role in metabolite production ^[7]^. Metabolic alterations during a critical window of development such as the fetal or immediate neonatal period are known to elicit life altering effects ^[8,9]^. Previous studies in adults show that antibiotic treatment disrupts intestinal homeostasis and has a profound impact on the intestinal metabolome ^[10]^. Antibiotics affect most metabolites that are detectable in the human intestinal tract, including ones critical for host physiology. Data on the effects of antibiotic use on the intestinal metabolome in preterm infants remains sparse.

Our group, designed a pragmatic randomized controlled pilot study to evaluate safety and feasibility of not giving routine antibiotics ^[11]^ after birth to symptomatic preterm infants considered at low risk for infection (REASON). This trial was also designed to evaluate longitudinally the fecal microbiome and metabolome in these infants. This trial randomized preterm infants with symptoms of prematurity to receiving (routine practice) vs. not receiving antibiotics shortly after birth and obtained weekly stool samples. Preliminary results showed that antibiotic use alters the microbiome and metabolome, including metabolites related to the gut-brain axis^[12]^. Of particular interest, gamma aminobutyric acid (GABA), which is a major neurotransmitter was found to be affected by antibiotic use. In the metabolomics sub analysis within the REASON trial, relative abundance of *Veillonella* and *Bifidobacterium* in preterm babies’ feces were both significantly affected by the administration of antibiotics. *Veillonella* showed a positive association with GABA, suggesting an antibiotic-mediated perturbation of a metabolite known to affect the gut-brain axis^[12]^. These variances could be significant for the lifetime health of the individual since they are occurring during a very sensitive window for development ^[9]^.

Administration of various types of antibiotics has also been shown in multiple animal models to produce significant changes on both the microbiome and the metabolome ^[13,14]^. One neonatal model theorized a personalized metabolome and microbiome based on antibiotic exposure ^[15]^. Here we describe the differences in fecal metabolomic profiles of preterm infants who were randomized to receive antibiotics vs no antibiotics after birth.

## 2. Results

### Study population

Neonates, born at gestational ages less than 33 weeks were sorted into 3 groups (A, C1, C2). Infants with symptoms not expected for gestation or infants at high risk for infection (Group B Streptococcus positive mother without adequate prophylaxis or chorioamnionitis) were placed into group A. Group C consisted of babies with symptoms expected for prematurity such as respiratory distress, and without significant risk factors for infection. Infants in group C were randomized to receive (C1(+) or not receive (C2(−)) antibiotics as described in Ruoss, et al.^[11]^ 3 infants from this subset were randomized to C2 (−) and changed to receive antibiotics in the 48hours after birth and therefore are included in the C1 (+) group for this analysis. 123 stools samples were collected: 43 for group A, 54 for group C1(+), and 26 for group C2(−).

### Metabolomics - comparison of 3 groups

PLS-DA cluster analysis of the metabolites (Figure 1A) depicts variances in the metabolic profiles between the three groups (Q2 = 0.49). The groups receiving antibiotics (A and C1(+) notably have more overlap in their metabolic profile while the no antibiotic group (C2(−)), although with some overlap, clusters further apart. 3D analysis of the randomized group C1 and C2 (Figure 1B) better illustrates the separation of their metabolites. The low Q2-value suggests there is more variability within the profiles and that more information would be necessary to for a more accurate model.

**Figure1.**
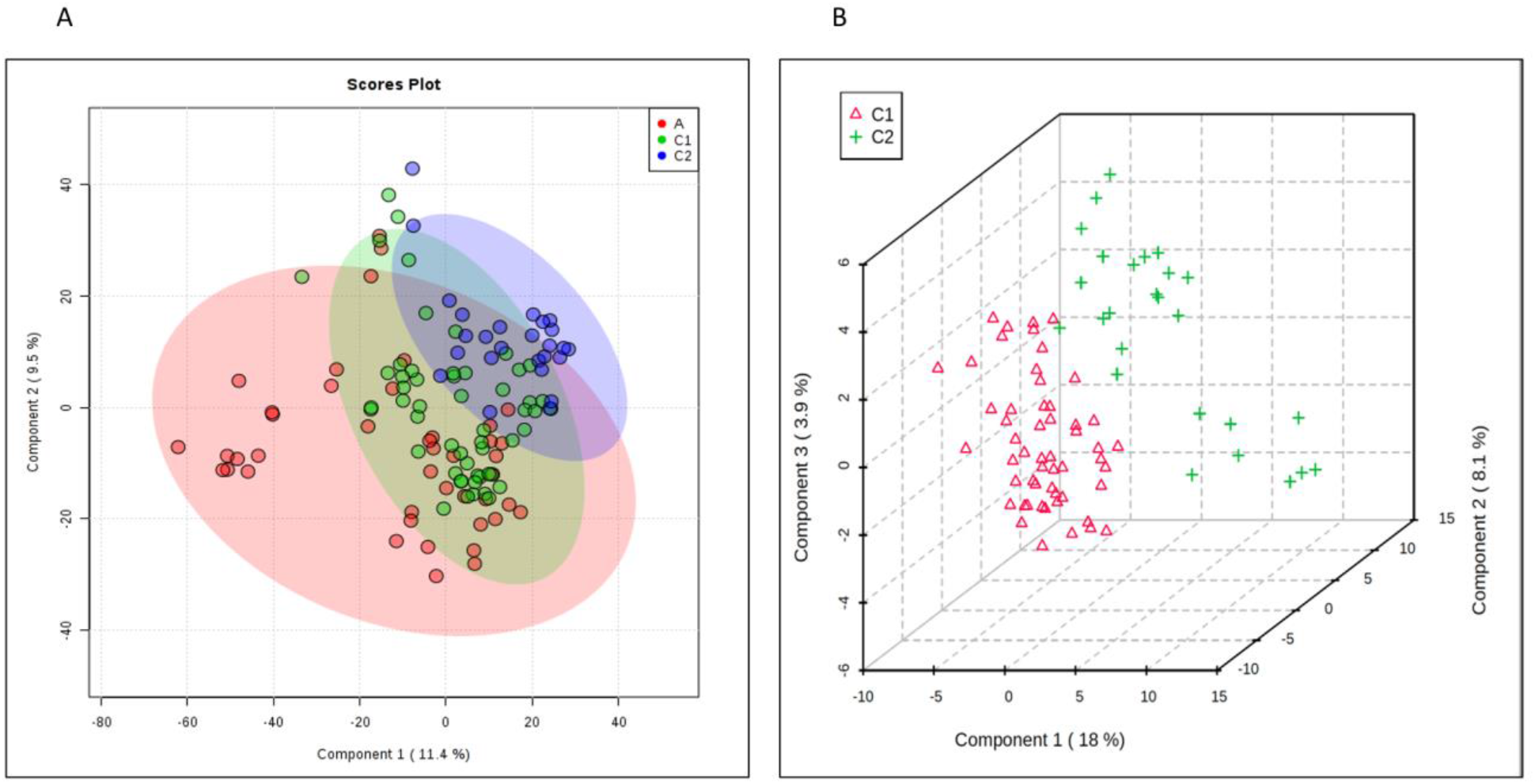
Comparison of Groups A, C1 and C2. **A.** PLS-DA plot depicting overlapping components of groups A (red), C1 (green) and C2 (blue). **B.** 3Dplot showing further delineation of groups C1 (red) and C2 (green).

Analysis using a volcano plot (Figure 2) demonstrates a few metabolites that are more significantly isolated between the groups, specifically gamma aminobutyric acid (GABA). This neurotransmitter correlated to certain microbial taxa, as described previously^[12]^.

**Figure 2.**
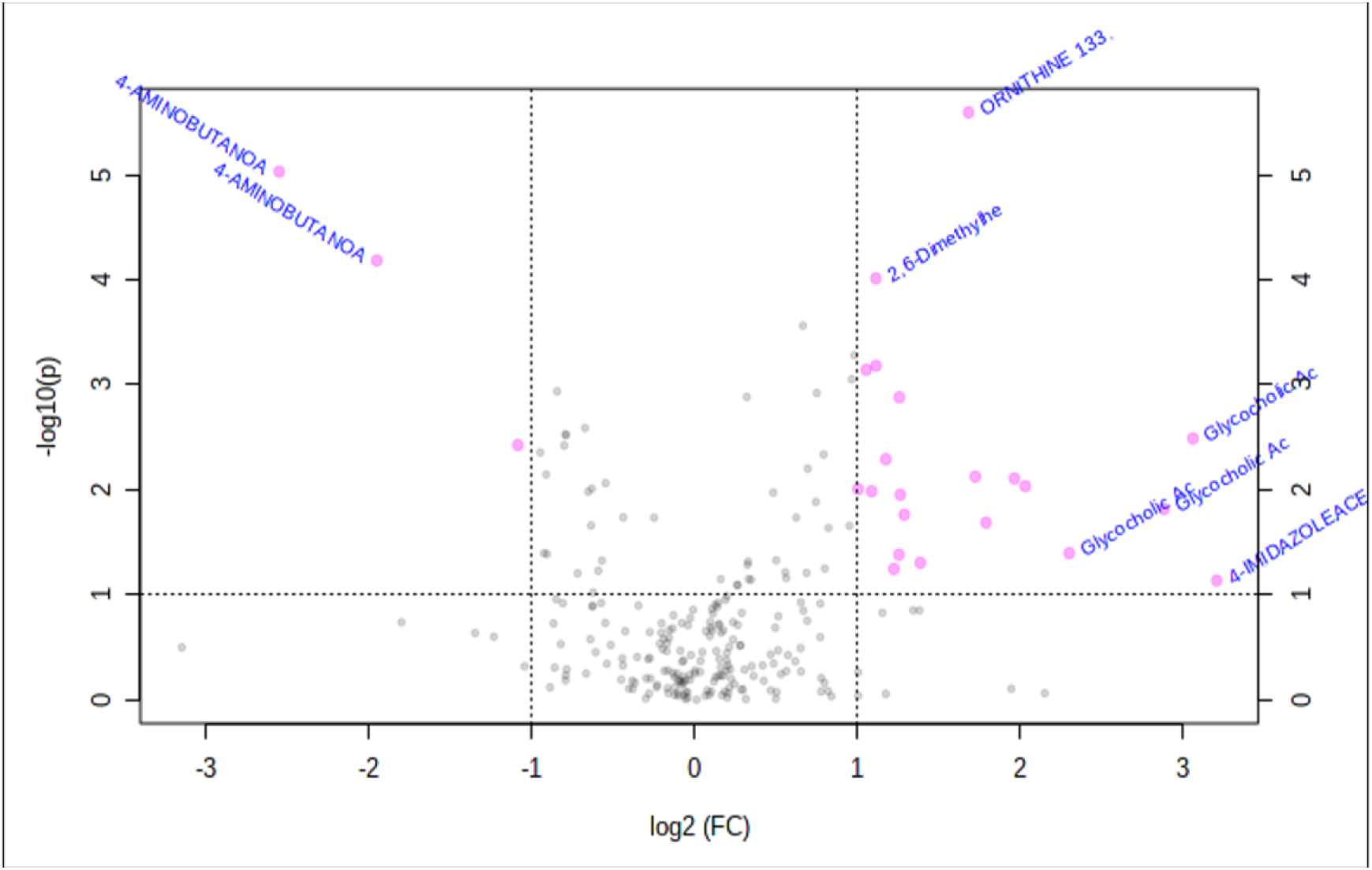
Volcano Plot. Illustrates the scatter of metabolites with increased significance at the extremes of the plot.

### Metabolomics—comparison of randomized groups

As differences observed in the PLS-DA and volcano plots differed, individual metabolites were compared. Here we will focus on several differences between within the infants randomized to receive or not to receive antibiotics (C1 vs. C2). The top 25 positive ion and 25 negative ion metabolites isolated within these samples are depicted via a heat map in Figure 3. It is apparent on these heat maps that the antibiotic group (red) and the no antibiotic group (green) not only have differences between them as denoted by the darkening red or blue colors, but also these appear to encompass categories of metabolites that will be discussed below.

**Figure 3.**
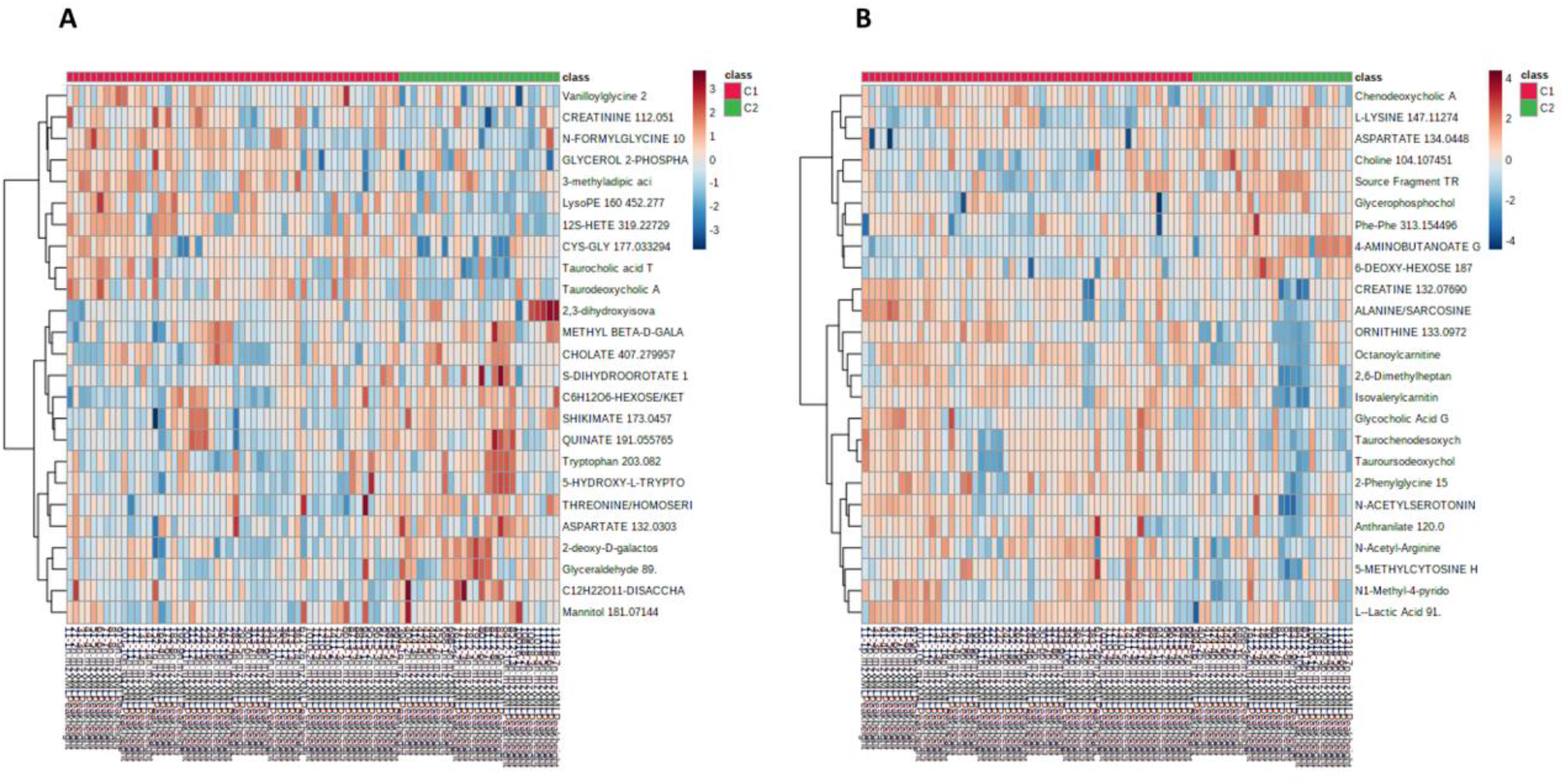
Heat Map. Color gradients depicting increased (red) versus decreased (blue) concentrations. **A.** Top 25 negative ion metabolites. **. B.** Top 25 positive ion metabolites.

Individual metabolites were compared using T-tests. The metabolites exhibiting significant (p<0.05) differences or intriguing trends are described in Table 1, where we also provide their function and group them into functional guilds. Figures 4–7 show these groupings.

**Table 1.**
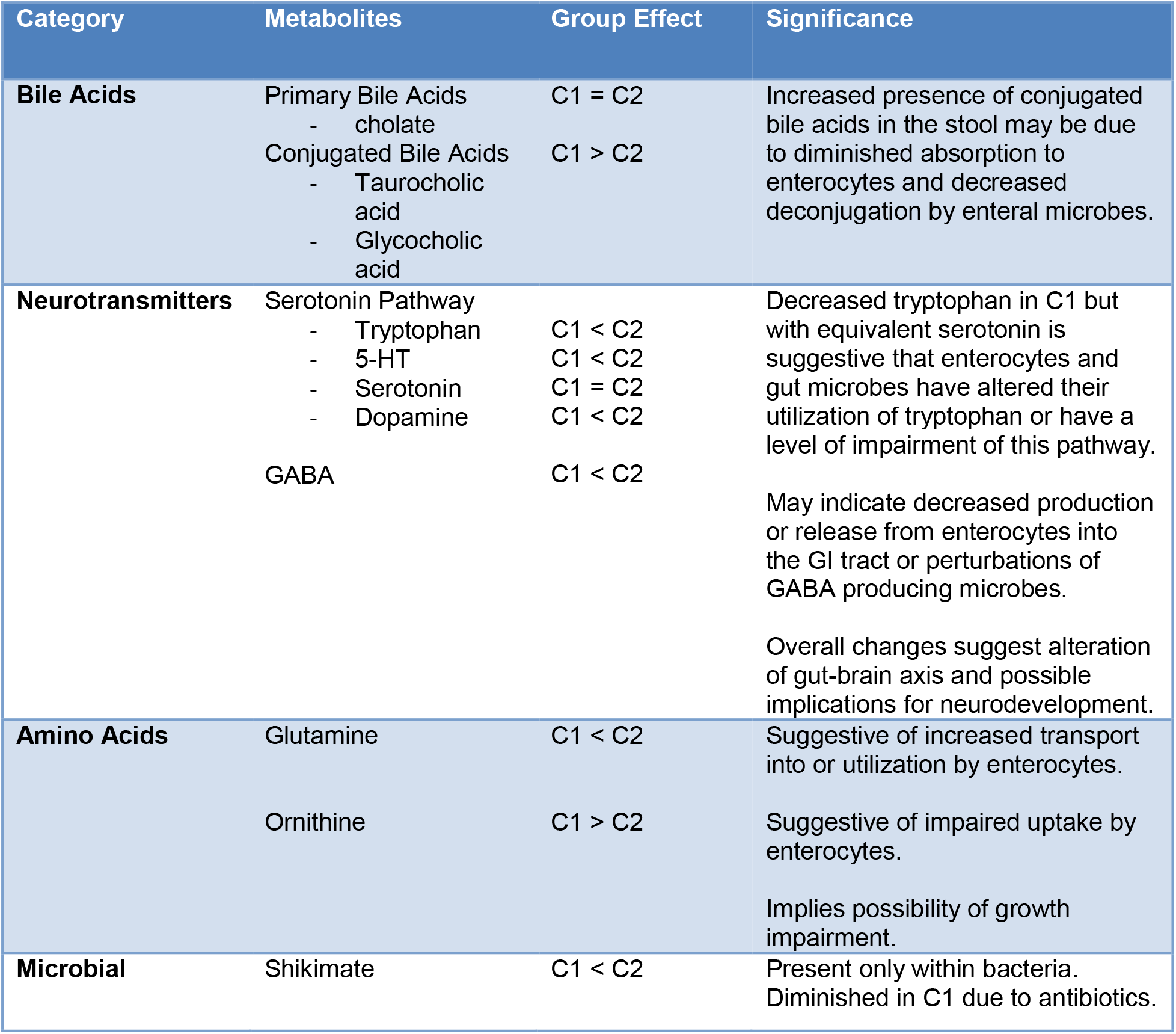
Classification, Summary, and Interpretation of Select Fecal Metabolites.

| Category | Metabolites | Group Effect | Significance |
| --- | --- | --- | --- |
| **Bile Acids** | Primary Bile Acids<br>- cholate | C1 = C2 | Increased presence of conjugated bile acids in the stool may be due to diminished absorption to enterocytes and decreased deconjugation by enteral microbes. |
|  | Conjugated Bile Acids<br>- Taurocholic acid<br>- Glycocholic acid | C1 > C2 |  |
| **Neurotransmitters** | Serotonin Pathway<br>- Tryptophan | C1 < C2 | Decreased tryptophan in C1 but with equivalent serotonin is suggestive that enterocytes and gut microbes have altered their utilization of tryptophan or have a level of impairment of this pathway. |
|  | - 5-HT | C1 < C2 |  |
|  | - Serotonin | C1 = C2 |  |
|  | - Dopamine | C1 < C2 |  |
|  | GABA | C1 < C2 | May indicate decreased production or release from enterocytes into the GI tract or perturbations of GABA producing microbes. |
|  |  |  | Overall changes suggest alteration of gut-brain axis and possible implications for neurodevelopment. |
| **Amino Acids** | Glutamine | C1 < C2 | Suggestive of increased transport into or utilization by enterocytes. |
|  | Ornithine | C1 > C2 | Suggestive of impaired uptake by enterocytes.<br>Implies possibility of growth impairment. |
| **Microbial** | Shikimate | C1 < C2 | Present only within bacteria. Diminished in C1 due to antibiotics. |

**Figure 4.**
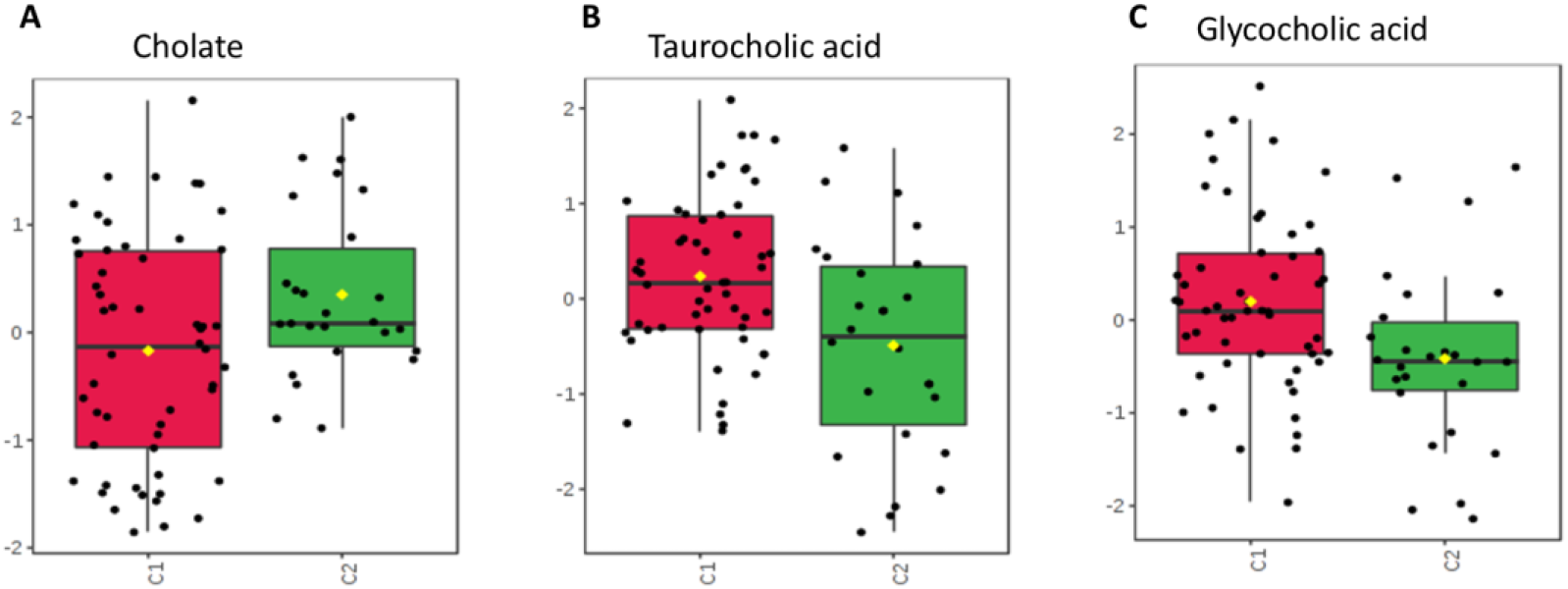
Concentrations compared between groups C1 and C2. **A.** Primary bile acid, cholate, with unchanged level in spite of antibiotic administration. **B** and **C**. Conjugated bile acids taurocholic acid and glycocholic acid. Both with increased expression in the group given antibiotics.

### Bile acid

Within the bile acid pathway, primary bile acids such as cholate and conjugated bile acids, such as taurocholic acid and glycocholic acid, figured more prominently within stool collected from the antibiotic group. No significant difference was seen in cholate level between the groups while both taurocholic acid (p=0.043) and glycocholic acid (p=0.047) were higher in antibiotic group comparing to no antibiotic group (Figure 4).

### Neurotransmitters

Neurotransmitters with significant physiologic activity within the gastrointestinal tract also varied between these groups (Figure 5). GABA demonstrated lower concentrations in fecal samples from the antibiotic group (p=0.001). Slight differences were also noted within the serotonin pathway, with the antibiotic group having a decreased fecal tryptophan (p=0.047) while seeming to maintain concentrations of 5-hydroxytryptamine (5-HT) (p=0.17), serotonin (p=0.82), and dopamine (p=0.40).

**Figure 5.**
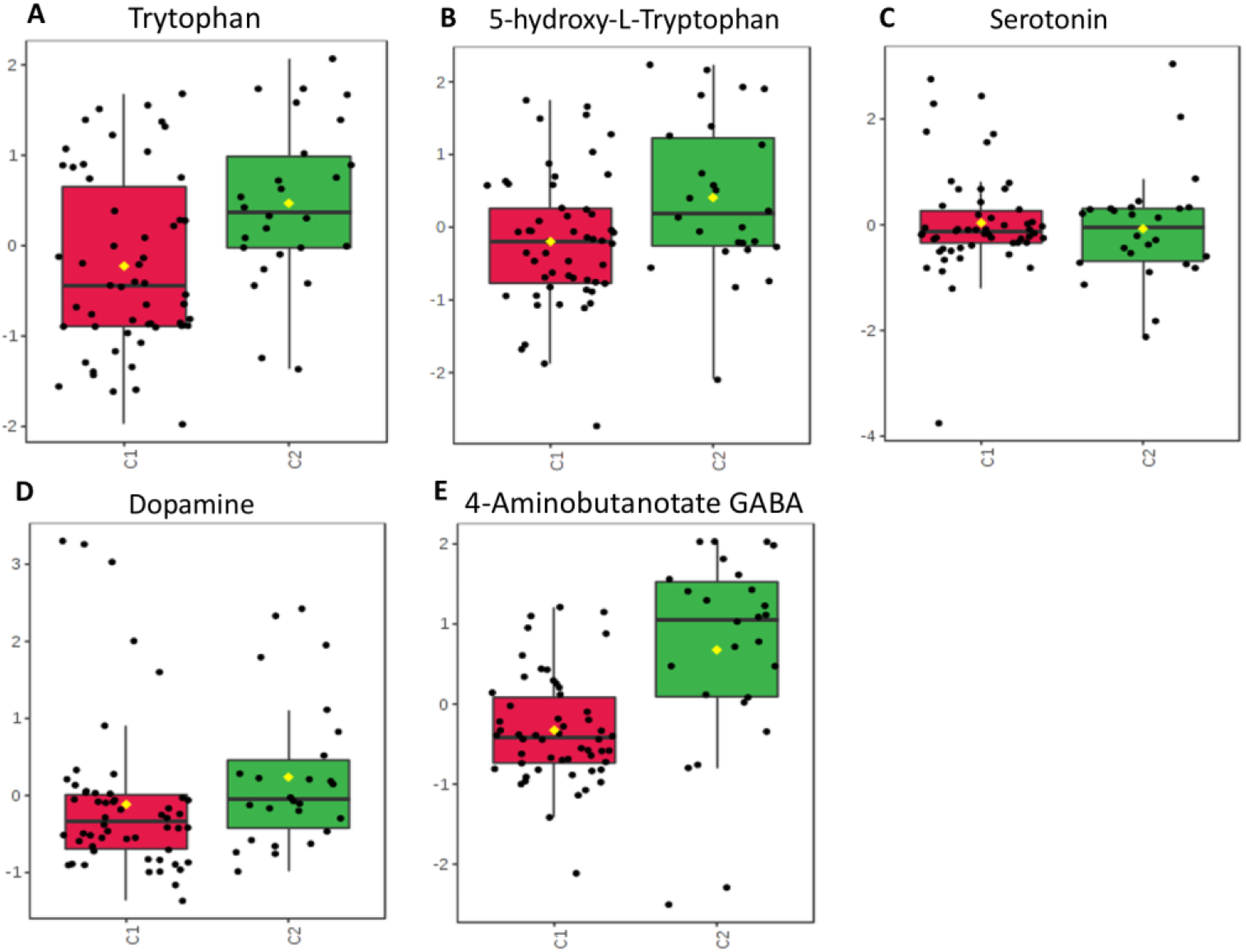
Neurotransmitters. Fecal concentrations compared between groups C1 and C2. **A-D**. Neurotransmitters falling within the serotonin pathway: tryptophan, 5-HT, serotonin, dopamine. **E.** GABA, a neurotransmitter which is produced in high quantities within enterocytes.

### Amino acids important to intestinal metabolism

Certain amino acids are highly utilized within the gastrointestinal tract both within microbes and enterocytes, some of which are involved with intestinal mucosal permeability (Figure 6). Of those isolated within these stools samples, ornithine had a lower fecal concentration within the no antibiotic group (p=0.001).

**Figure 6.**
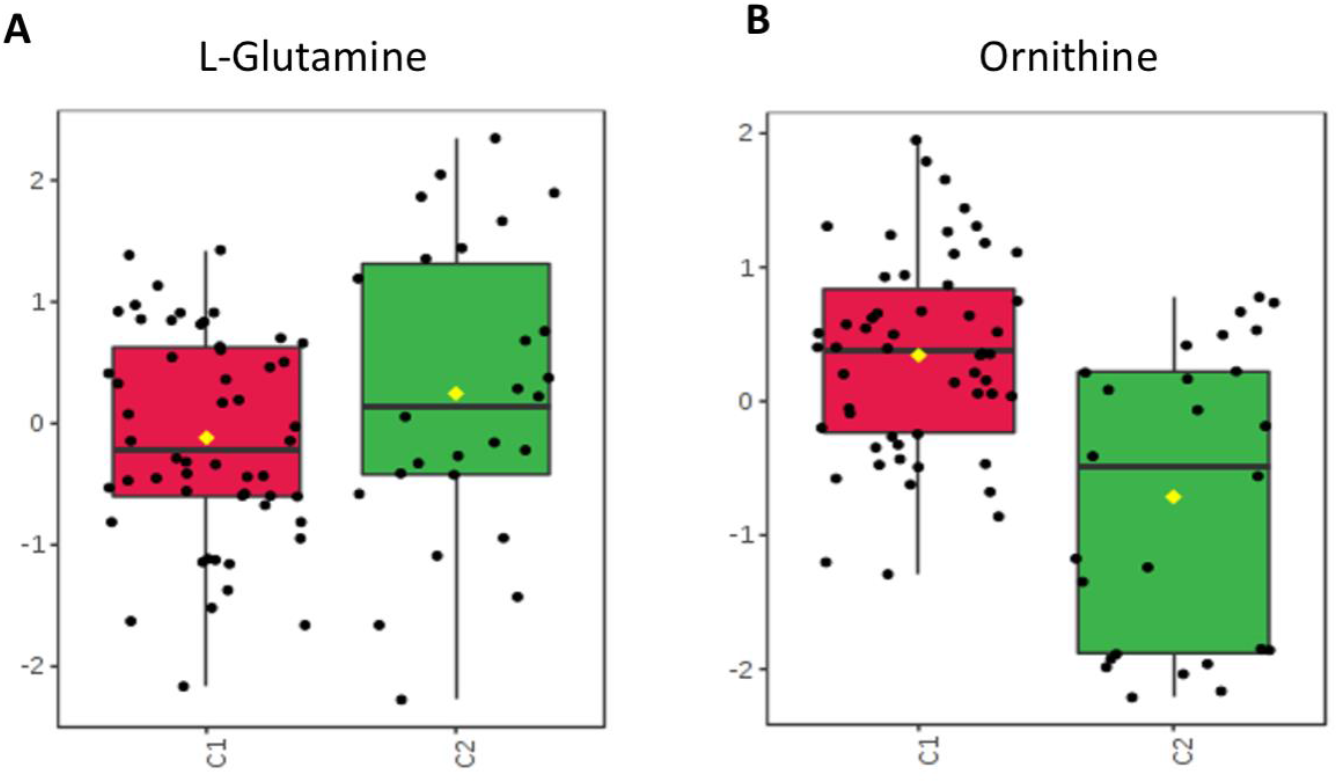
Amino Acids. Fecal concentrations compared between groups C1 and C2. **A.** Glutamine, with a trend of decreased concentration in the group exposed to antibiotics. **B.** Ornithine, with increased concentration in the group exposed to antibiotics.

### Shikimate-Folic acid metabolism

Shikimate, utilized by bacteria in the production of folates and aromatic amino acids, was noted in higher concentrations within the no antibiotic group comparing to antibiotic group (Figure 7, p=0.043). As humans are not able to utilize or process these metabolites, alterations in their concentration would result from exposure or changes within the gastrointestinal environment.

**Figure 7.**
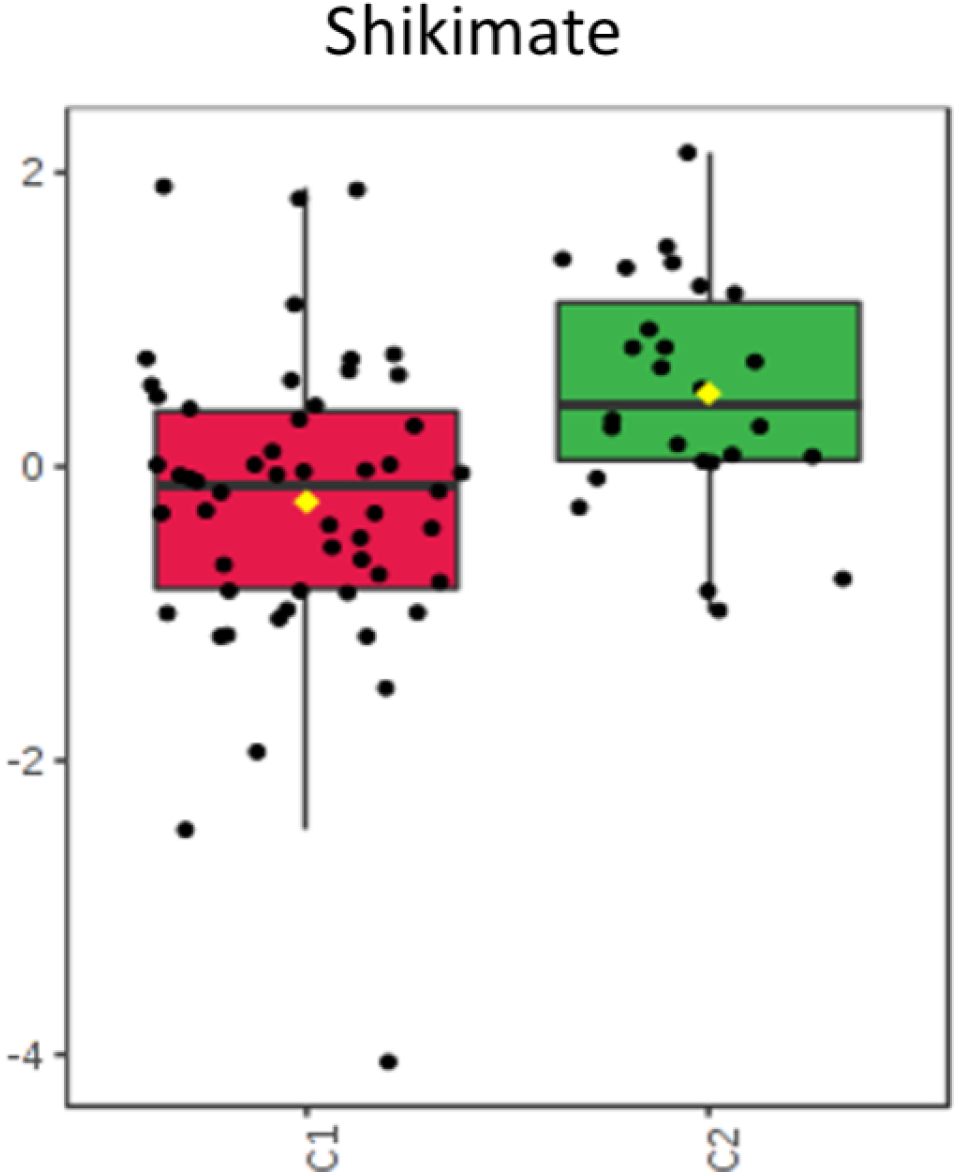
Concentrations compared between groups C1 and C2. Shikimate, bacterial metabolite important within the shikimate-folate pathway.

## Discussion

Antibiotic administration within the NICU is a common practice given concerns for sepsis around the time of delivery when few clinically measurable factors are reliable indicators of neonatal infection. Exposure to antibiotics early in life alters the developing microbiome thereby affects the metabolome. The gastrointestinal tract plays significant roles in terms of nutrition, immunity, and even neurotransmitter production. Alterations in the microbiome and metabolome using antibiotics could then significantly impair or alter these functions and predispose those babies to future complications. Studies analyzing fecal metabolites published so far have been non-randomized, observational, or retrospective. This study is prospective and randomized, aiming to identify and evaluate metabolites within the stool of preterm neonates specifically related to early antibiotic administration.

Of the isolated metabolites compared between the antibiotic and no antibiotic groups, some show more significant differences and warrant further consideration. These are categorized as follow:

### Bile acids

Some of the more prominently portrayed and variant metabolites are bile acids (Figure 4). The neonates receiving antibiotics showed a higher expression of conjugated bile acids (glycocholic acid and taurocholic acid). These are typically reabsorbed within distal portions of the gastrointestinal tract^[16]^ and this increase may be secondary to decreased absorption into intestinal cells. Some intestinal microbes, such as Bifidobacteriae, are also capable of deconjugating these bile acids back into primary acids^[17,18]^, and Russell et al. noted a negative correlation between the abundance of Bifidobacteriae and the conjugated bile acids between neonates given antibiotics and those without antibiotics^[12]^. As antibiotic administration changes the intestinal microbiome, bacteria capable of converting these conjugated bile acids may be significantly reduced or completely gone. This then would lead to increased excretion of glycocholic acid and taurocholic acid and could also explain the diminished expression of cholate in the antibiotic group. Bile acids are important for lipid absorption within the intestines ^[19]^ so changes seen between these groups could signify abnormalities of nutrient exchange with implications for nutrition and growth of these neonates.

### Neurotransmitters

Tryptophan, serotonin, dopamine, and GABA are all neurotransmitters that are interestingly found in high quantities within the gastrointestinal tract. Our results show alterations in fecal expression of these neurotransmitters between groups randomized with and without antibiotics. (Figure 5) The lower concentration of tryptophan is noted within the group receiving antibiotics while serotonin appears unchanged. Tryptophan is an essential amino acid, and as such, its presence within the gut is modulated by diet ^[20]^. The gut is also the main source of serotonin synthesis, with over 90% of serotonin being produced by enterochromaffin cells, mucosal mast cells, and myenteric cells ^[21]^. Some microbes, such as Pseudomonas, are capable of converting tryptophan into serotonin for their own metabolism ^[20]^. If antibiotics allow colonies of such microbes to grow, fecal tryptophan levels would diminish while fecal serotonin would remain stable. The decreased availability of tryptophan to enterocytes may subsequently decrease the amount of serotonin available for systemic distribution. Conversely, if antibiotics are decreasing the variety of microbes within the GI tract, the enterocytes may have increased access to and utilization of tryptophan to further process to serotonin. As fecal serotonin is seemingly unchanged between these groups, this short course of antibiotics may have diminished bacterial levels to either allow formation of serotonin producing microbes or allowed for increased utilization by enterocytes.

GABA was also found in lower concentrations among the group randomized to receive antibiotics. Produced within the myenteric plexus and within mucosal endocrine cells, GABA plays a role in contraction and relaxation of enteric muscles and may also play a role in gastrointestinal immunity ^[22]^. As a neurotransmitter and part of the gut-brain axis, GABA is important for neurodevelopment ^[23]^. Russell et al. showed in this preterm population that GABA concentrations were negatively impacted by the administration of antibiotics at least partly due to decreased colonization with *Veillonella*^[12]^. Lactobacillus and Bifidobacteria have also been shown to produce GABA ^[24]^. Realistically, changes in the microbiota from antibiotics could lead to significant alterations in GABA producing bacteria that may contribute to our results. Given our developing understanding of the gut-brain axis and GABA’s role in neurodevelopment, as well as its role in immunity, decreased levels of GABA in stool may imply larger effect on neonatal development.

### Amino acids critical to Intestinal metabolism

Ornithine and glutamine are amino acids highly utilized within the healthy gastrointestinal tract. Glutamine provides an energy source for enterocytes and aids with protein synthesis as well as differentiation of enterocytes ^[25]^. Models of sepsis and gut inflammation have shown decreased uptake of glutamine by enterocytes^[25]^ and supplementation of glutamine has been studied in prevention of NEC in preterm infants as it has been found to decrease the possibility of bacterial translocation across enterocytes^[26]^. The decreased concentration in the antibiotic group may be secondary to increased utilization in maintenance of the mucosal barrier and in further proliferation of enterocytes.(Figure 6) This response may be from a reaction within the enterocytes due to microbial changes caused from the short course of antibiotics and may demonstrate degradation of metabolic and immune function of these enterocytes. Similarly, enterocytes actively transport and utilize ornithine as an important energy source and it is important for appropriate growth ^[27]^ as it can stimulate the release of growth hormone^[28]^. Elevated levels in the stool of the antibiotic group suggest that absorption into enterocytes likely is impaired. This disruption could impact the overall growth and subsequent development of a neonate, even though these antibiotics were only administered for a very short period of time.

### Shikimate

Shikimate is a metabolite whose presence within the fecal metabolome is completely dependent on production by intestinal microbes as humans are not capable of producing it^[29]^. It is noted within both groups but has a larger concentration within the no antibiotic group.(Figure 7) Alterations in its expression are associated with antibiotic use and suggest that even a short course of antibiotics can produce a significant change in these important metabolites.

## Methods

### Clinical trial

The REASON study was conducted from January 2017 - January 2019 at the University of Florida after approval by the institutional review board (IRB201501045). A detailed description of the study design including enrollment, group selection, randomization, and collection of clinical samples and data including outcomes has been previously described ^[11]^. Briefly, 98 infants <33 weeks gestation were enrolled and placed into one of three groups according to previously described criteria: group A neonates with high risk for infection with indication for antibiotic treatment, group B asymptomatic babies with low risk for infection without indication for antibiotics, and group C symptomatic babies with low risk for infection, eligible for randomization to antibiotics (group C1) or no antibiotics (group C2). Infants in group C1 were placed on antibiotics for 48 hours after birth. Infants could be continued on antibiotics longer then 48hours in group C1 or started on antibiotics within 48 hours after birth in group C2 based on the medical team’s judgement. Because of financial limitations only the first 19 patients, including group A (n=6, with 43 weekly samples), group C1 (n=8, with 54 weekly samples) and group C2 (n=5, with 26 weekly samples), analyzed in the REASON trial^[11]^ underwent fecal metabolite analysis and are presented in this paper.

### Stool sample collection

Weekly fecal samples, including first meconium when possible, were collected and stored at −80°C. The infant stool samples were suspended in 400 μL 5 mM Ammonium Acetate. Homogenization was done 3 times for 30 s each time using a cell disruptor. Protein concentrations of the homogenates were measured using Qubit Protein Assay. Samples were normalized to 500 μg/mL protein at 25 μL for extraction. Each normalized sample was spiked with 5 μL of internal standards solution. Extraction of metabolites was performed by protein precipitation by adding 200 μL of extraction solution consisting of 8:1:1 Acetonitrile: Methanol: Acetone to each sample. Samples were mixed thoroughly, incubated at 4°C to allow protein precipitation, and centrifuged at 20,000 × G to pellet the protein. 190 μl supernatant was transferred into clean tube and dried using nitrogen. Samples were reconstituted with 25 μL of reconstitution solution consisting of injection standards, mixed, and incubated at 4°C for 10-15 min. Samples were centrifuged at 20000 × G. Supernatants were transferred into LC-vials.

### Metabolomic profiling

Global metabolomics profiling was performed on a Thermo Q-Exactive Orbitrap mass spectrometer with Dionex UHPLC and autosampler ^[30]^. All samples were analyzed in positive and negative heated electrospray ionization with a mass resolution of 35,000 at *m/z* 200 as separate injections. Separation was achieved on an ACE 18-pfp 100 × 2.1 mm, 2 μm column with mobile phase A as 0.1% formic acid in water and mobile phase B as acetonitrile. The flow rate was 350 μL/min with a column temperature of 25°C. 4 μL was injected for negative ions and 2 μL for positive ions.

Data from positive and negative ions modes were processed separately. LC-MS files were first converted to open-format files (i.e. mzXML) using MSConvert from Proteowizard. Mzmine was used to identify features, deisotope, align features and perform gap filling to fill in any features that may have been missed in the first alignment algorithm. Features were matched with SECIM internal compound database to identify metabolites ^[31]^. All adducts and complexes were identified and removed from the data set prior to statistical analysis.

### Statistical Analysis

Metaboanalyst 3.0 was used for all statistical analysis including PCA, volcano plot, ANOVA, T tests, heatmaps analysis and individual comparison of the metabolites that exhibited the highest significance as determined by p-value less than or equal to 0.05^[32]^.

## Limitations and Future Directions

Despite a limited sample size, several significant associations were found in fecal metabolites based on antibiotic use. In future studies, a larger sample size will allow for analysis of these metabolites throughout the course of each neonate’s hospitalization and in response to individual antibiotic courses.

Across these samples we have shown a variety of isolated metabolites, although only a small proportion of these are currently identifiable. As more information is collected through all research involving metabolites, the national database that we use for purposes of identification will grow and more of these unknowns can be categorized and utilized for theories of importance. Despite these limitations, the antibiotic associated alterations in metabolites seen during the critical developmental window of preterm infants prompts the need for a more in-depth study, the results of which could be considered when the decision is made to routinely provide antibiotics without clear evidence of infection.

## Acknowledgements

This study was funded by the NIH under grant R21HD088005.

## Author contributions

Conceptualization, Josef Neu; Data curation, Laura Patton, Timothy J. Garrett and J. Lauren Ruoss; Formal analysis, Nan Li and Timothy J. Garrett; Funding acquisition, Josef Neu; Investigation, Laura Patton, Nan Li, Timothy J. Garrett, J. Lauren Ruoss, Jordan T. Russell, Diomel de la Cruz, Catalina Bazacliu and Josef Neu; Methodology, Nan Li, Timothy J. Garrett and J. Lauren Ruoss; Project administration, Josef Neu; Supervision, Josef Neu; Validation, Laura Patton, Timothy J. Garrett, Richard A. Polin, Eric W. Triplett and Josef Neu; Writing – original draft, Laura Patton and Nan Li; Writing – review & editing, Timothy J. Garrett, J. Lauren Ruoss, Jordan T. Russell, Diomel de la Cruz, Catalina Bazacliu and Josef Neu.

